# Inhibition of SARS-CoV-2 in Vero cell cultures by peptide-conjugated morpholino-oligomers

**DOI:** 10.1101/2020.09.29.319731

**Authors:** Kyle Rosenke, Shanna Leventhal, Hong M. Moulton, Susan Hatlevig, David Hawman, Heinz Feldmann, David A. Stein

## Abstract

**Background:** SARS-CoV-2 is the causative agent of COVID-19 and a pathogen of immense global public health importance. Development of innovative direct-acting antiviral agents is sorely needed to address this virus. Peptide-conjugated morpholino oligomers (PPMO) are antisense agents composed of a phosphordiamidate morpholino oligomer covalently conjugated to a cell-penetrating peptide. PPMO require no delivery assistance to enter cells and are able to reduce expression of targeted RNA through sequence-specific steric blocking.

**Objectives and Methods:** Five PPMO designed against sequences of genomic RNA in the SARS-CoV-2 5’-untranslated region and a negative control PPMO of random sequence were synthesized. Each PPMO was evaluated for its effect on the viability of uninfected cells and its inhibitory effect on the replication of SARS-CoV-2 in Vero-E6 cell cultures. Cell viability was evaluated with an ATP-based method and viral growth was measured with quantitative RT-PCR and TCID_50_ infectivity assays.

**Results:** PPMO designed to base-pair with sequence in the 5’-terminal region or the leader transcription regulatory sequence-region of SARS-CoV-2 genomic RNA were highly efficacious, reducing viral titers by up to 4-6 log10 in cell cultures at 48-72 hours post-infection, in a non-toxic and dose-responsive manner.

**Conclusion:** The data indicate that PPMO have the ability to potently and specifically suppress SARS-CoV-2 growth and are promising candidates for further pre-clinical development.

## Introduction

There is a pressing need for the development of additional antiviral therapeutics to address SARS-CoV-2 infections. Peptide-conjugated phosphorodiamidate morpholino oligomers (PPMO) are single-stranded-nucleic acid analogs capable of entering cells without assistance and interfering with gene expression through steric blockade of targeted RNA. PPMO are composed of a phosphorodiamidate morpholino oligomer (PMO) covalently conjugated to a cell-penetrating peptide (CPP)[1–3]. PPMO are water-soluble, nuclease resistant and non-toxic at effective concentrations across a range of *in vitro* and *in vivo* applications [1, 4]. In several *in vivo* models of respiratory viruses, including influenza A virus[5, 6], respiratory syncytial virus [7]and porcine reproductive and respiratory syndrome virus[8], intranasally administered PPMO targeted against virus sequence have reduced viral titer and pathology in lung tissue.

The 5’UTR of the coronavirus genome contains sequences and structures known to be important in various aspects of the virus life-cycle including translation and RNA synthesis [9]. In previous studies, PPMO targeted to various sites in the 5’UTR of mouse hepatitis virus (MHV) [10, 11] and SARS-CoV [12] were effectively antiviral. In the present study we investigated the ability of five PPMO targeted against various sites in the 5’UTR and polyprotein 1a/b translation start site (AUG) region of SARS-CoV-2 to suppress virus growth in cell cultures. We found that PPMO targeting the 5’terminal region and the transcription regulatory sequence (TRS)-leader region of genomic RNA were potent inhibitors of SARS-CoV-2 replication.

## Materials and Methods

### Biosafety

Work with infectious SARS-CoV-2 was approved by the Institutional Biosafety Committee (IBC) and performed in high biocontainment at Rocky Mountain Laboratories (RML), NIAID, NIH. Sample removal from high biocontainment followed IBC-approved Standard Operating Protocols.

### PPMO synthesis

PPMO were synthesized by covalently conjugating the CPP (RXR)4 (where R is arginine and X is 6-aminohexanoic acid) to PMO (Gene Tools LLC, Philomath, Oregon) at the 3’ end through a noncleavable linker, by methods described previously [3].

### Cells and viruses

Vero-E6 cells (ATCC) were maintained in DMEM supplemented with 10% fetal calf serum, 1 mM L-glutamine, 50 U/mL penicillin and 50 μg/mL streptomycin (growth medium). All cell culture incubations were carried out at 37° C and 5% CO_2_. SARS-CoV-2 isolate nCoV-WA1-2020 was kindly provided by the Centers for Disease Control and Prevention (Atlanta, Georgia, USA). Preparation and quantification of the virus followed methods previously described[13]. Briefly, the original virus stock was propagated once at RML in Vero-E6 cells in DMEM supplemented with 2% fetal bovine serum containing L-glutamine and antibiotics as above (infection medium). The virus stock used in the experiments was passage 4 and was confirmed by sequencing to be identical to the initial deposited GenBank sequence MN985325.

### Cell viability assay

Cell viability was assessed using CellTiterGlo (Promega). Vero E6 cells grown in 96-well plates were incubated in the presence of PPMO in growth media for 48 hours, then assayed according to the manufacturer’s instructions. Statistical analysis was carried out using GraphPad Prizm (San Diego, CA).

### Antiviral assays

PPMO were resuspended in sterile PBS. Vero-E6 cells were plated in 48 well plates and were approximately 80% confluent on the day of infection. At 5 hrs before infection, the growth medium was removed and replaced with infection medium containing PPMO. For viral infections, the PPMO-containing medium was aspirated and the cells rinsed twice with DMEM before adding 100 μl of infection medium containing virus at a MOI of .01. Following a one hour infection period, the virus-containing inoculum was aspirated and the cells washed twice with infection medium after which 300 μl fresh infection medium was added. At 12, 24, 48 and 72 hours post-infection (pi), all of the media in a well was collected for qRT-PCR and TCID_50_ analysis.

### Evaluation of virus quantity by qRT-PCR

Supernatants were harvested as described above and viral RNA purified and quantified by using one-step quantitative reverse transcription PCR (qRT-PCR) following previous described methods [13]. Briefly, total RNA was isolated with the RNeasy Mini kit (Qiagen) and RT-PCR carried out using the One-Step RT-PCR kit (Qiagen) according to the manufacturer’s protocols. Copy numbers were calculated using standards produced as previously described [13].

### TCID_50_ evaluations

Viral supernatants were serially diluted in DMEM and each dilution sample was titrated in triplicate. Subsequently, 100 ul of each virus dilution was transferred to Vero-E6 cells grown in 96-well plates containing 100ul DMEM. Following a seven day incubation period, wells were scored for cytopathic effect (CPE). The TCID_50_ values were calculated via the Reed-Muench formula.

## Results

### PPMO design

PPMO design was guided by previous studies in which various Nidoviruses were targeted with PPMO [10–12, 14, 15]. In this study, five PPMO were designed to target the 5’ UTR and first translation start site-region of SARS-CoV-2 positive sense genomic RNA (**Table 1**). Two of the PPMO target the 5’-terminal-region of the genome. Both 5’END-1 and 5’END-2 were designed with the intention of interfering with pre-initiation of translation of genomic and subgenomic mRNAs.

**Table 1.**
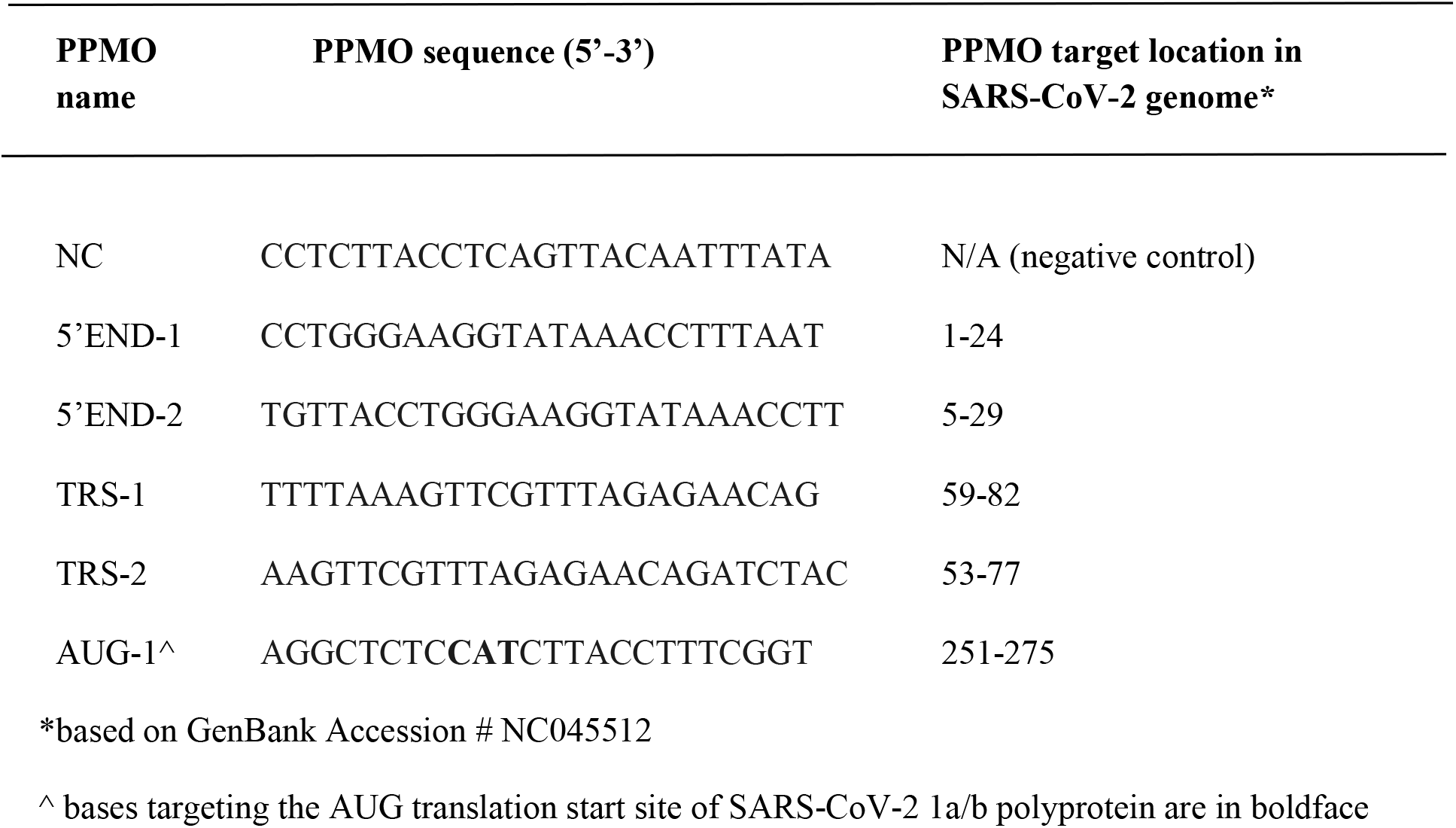
Names, sequences, and target locations of SARS-CoV-2 PPMO

Two PPMO were designed to target the genomic 5’UTR region containing the putative SARS-CoV-2 leader-TRS core sequence (nt 70-75, 5’-ACGAAC-3’) and thereby potentially interfere with body-TRS to leader-TRS base-pairing, and/or with translocation of the 48S translation preinitiation complex along the 5’UTR of the genomic and various subgenomic mRNAs. Coronaviruses produce a set of nested mRNAs through the process of discontinuous subgenomic mRNA synthesis. The TRS is a six-ten nt sequence that is critical in the production of negative strand mRNA templates during this process [16, 17]. For SARS-CoV, the leader-TRS core sequence consists of nt 67-72 of the genomic RNA sequence[18], and most viral mRNAs possess the same 72 nt 5’ leader sequence.

The AUG PPMO was designed to target the translation initiation region for ORF1a/b, which codes for the replicase polyprotein, with the intention to block translation initiation.

A negative control PPMO (NC) of random sequence was included (see Table 1), to control for nonspecific effects of the PPMO chemistry. NC was screened using BLAST and contains little significant homology to any primate, rodent or viral sequences.

### Evaluation of PPMO cytotoxicity

To evaluate the effect of PPMO treatments on cell viability, cells were treated under similar conditions to the antiviral assays described below, but in the absence of virus. Cells were treated in triplicate with increasing doses of PPMO for 48 hours before being assayed using a quantitative cell viability assay. At the concentrations used in the antiviral assays described below, none of the PPMO produced more than 5% cytotoxic effect (**Figure 1A**).

**Figure 1.**
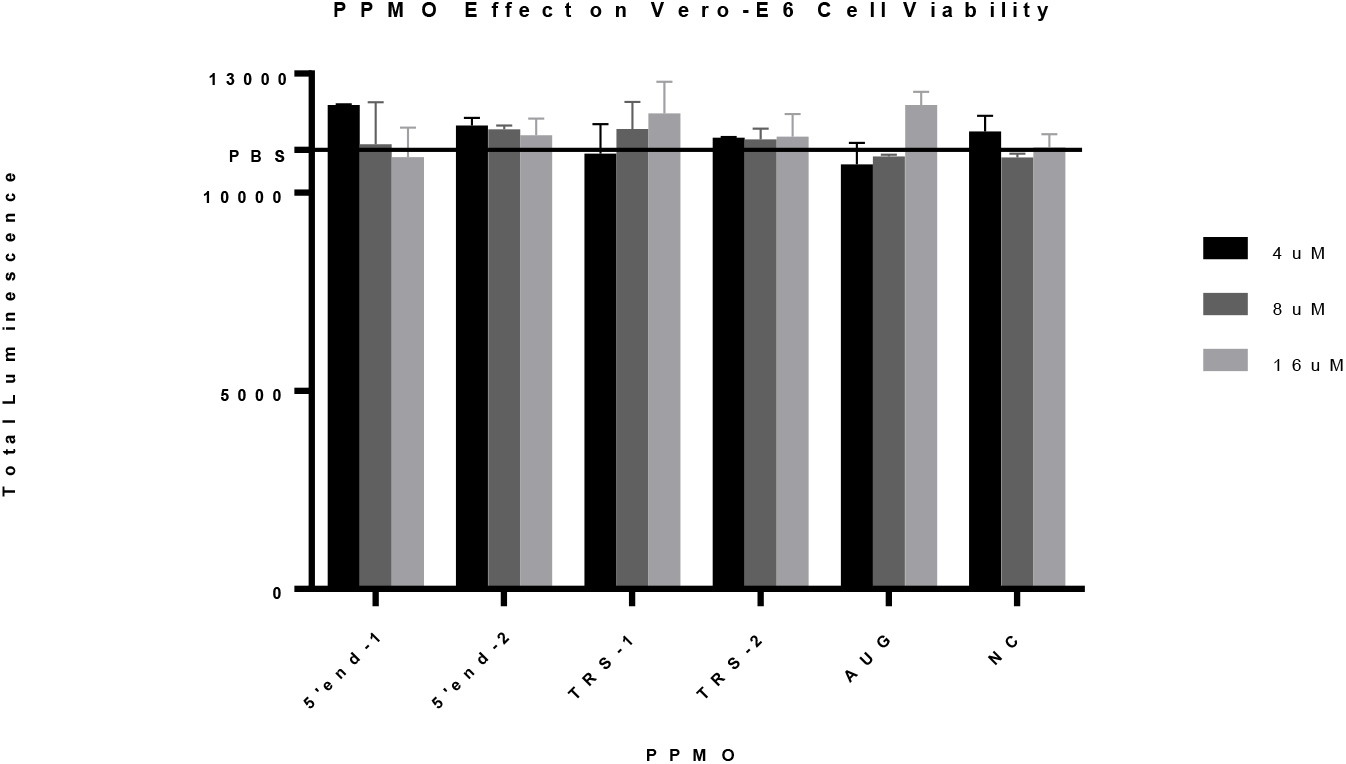
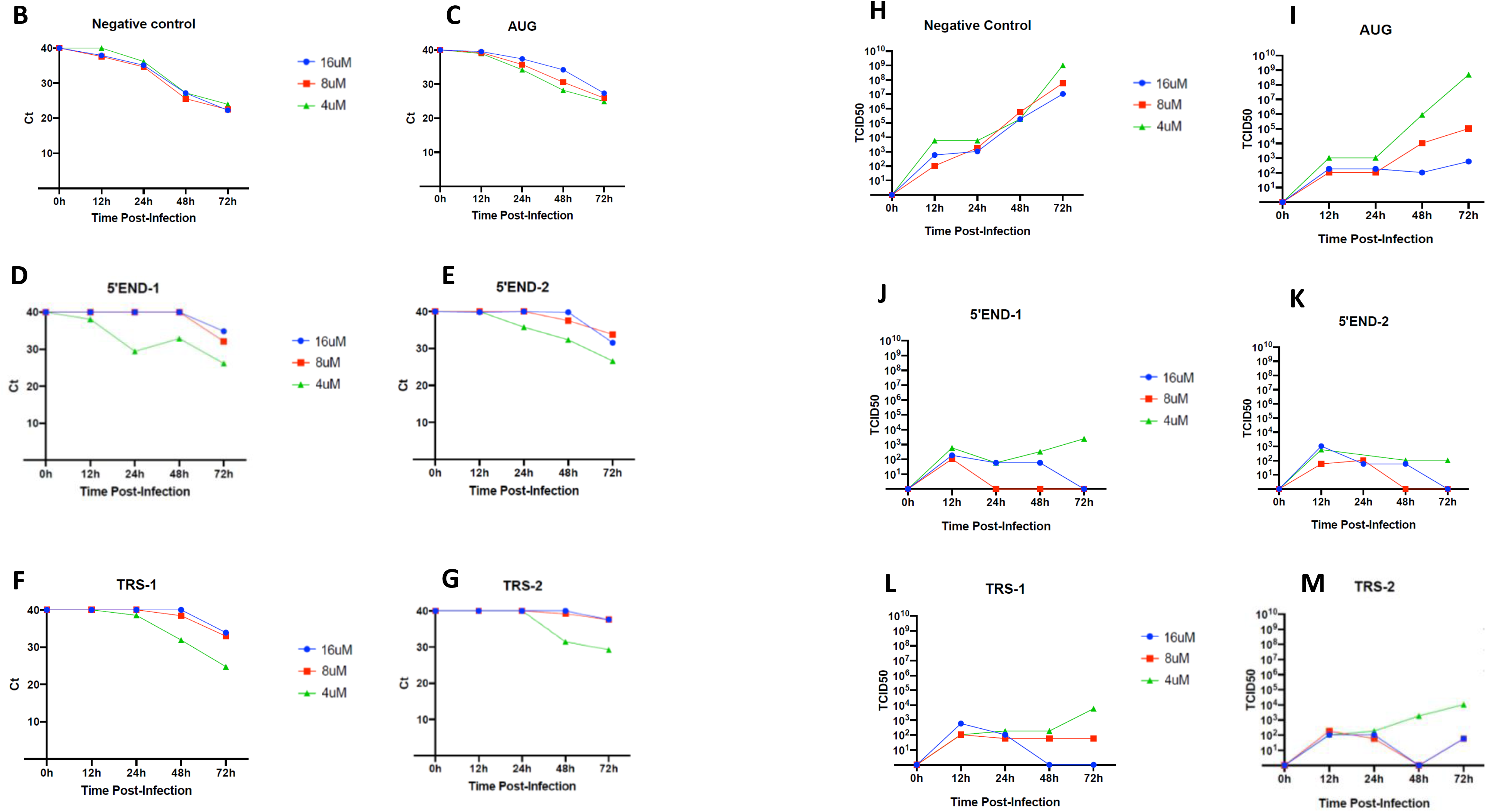
Effect of PPMO on cell viability and virus growth. **A**) the effect of PPMO treatment on cellular ATP level as an indicator of cell viability was carried out using uninfected cells incubated for 48 hours with increasing concentrations of the indicated PPMO. ATP levels were determined via luminescence readings and are shown compared to PBS-treated cells. For each PPMO concentration, triplicate samples were assayed and the mean +/- standard deviation is shown. **B-M**) Growth curves of SARS-CoV-2. Vero-E6 cells were treated with indicated concentration of PPMO for 5 hours before infection with MOI of .01 of SARS-CoV-2. Cell supernatants were collected at 12, 24, 48 and 72 hours post-infection and analyzed by quantitative RT-PCR (B-G) or TCID_50_/ml end-point dilution (H-M) with 3 technical repeats. Cells treated with PBS only had titers similar to NC PPMO. The limit of virus detection for the TCID_50_ assay is 10^1^ /ml. This experiment was carried out twice, under similar conditions, yielding similar results, and a single representative experiment is shown.

### Evaluation of PPMO antiviral activity

To determine the inhibitory activity of the various PPMO on SARS-CoV-2 replication, Vero-E6 cells were treated with each of the six PPMO described in Table 1 at 4, 8, and 16 μM for 5 hours before infection, then incubated without PPMO after infection. Cell supernatants were collected at four time-points post-infection: 12, 24, 48, and 72 hours. Virus growth was evaluated by two methods, qRT-PCR and TCID_50_ infectivity assay. Using an MOI of 0.01, virus growth rose steadily and reached peak growth at 72 hrs post-infection (**Fig. 1B and H**). Growth of the virus under PBS treatment was highly similar to virus growth under NC PPMO treatment (data not shown). Four of the five PPMO designed to target SARS-CoV-2 RNA were highly effective, suppressing viral titers by 3-5 log10 at the 48 and 72 hr time-points (**Figure 1B-M**). qPCR analysis showed that in cells treated with 8 or 16 μM of any of the four PPMO targeting the 5’end- or TRS-regions, virus growth was markedly suppressed at 12-48 hrs post-infection. The number of cycles required to detect virus from those samples was approximately the same as the number of cycles required to detect the input virus shortly after infection. At 72 hrs post-infection, the efficacy of the 5’END- and TRS-PPMO had waned, to a minor extent, although still providing multi-log suppression of virus growth. The AUG PPMO was not nearly as effective as the other 4 antiviral PPMO used in this study (**Figure 1C and I**).

## Discussion

We found that PPMO targeting the 5’terminal-region or leader-TRS-region were highly effective at inhibiting the growth of SARS-CoV-2, whereas a PPMO targeting the polyprotein 1a/b AUG translation start site region was not effective. It is unknown if the ineffective AUG-PPMO was unable to bind to its target, or if duplexing occurred yet was relatively inconsequential.

To date, little sequence variation in the PPMO target sites in the 5’UTR of SARS-CoV-2 has been identified. Of the whole-genome nucleotide sequences reported in GenBank, two genotypes contain a single mismatch with the 5’END PPMO and one genome has a single mismatch with the TRS PPMO[19]. Previous studies have shown that PPMO having a single base mismatch with their target site retain approximately 90% of their activity compared to those having perfect agreement [15, 20].

This study demonstrates that PPMO targeted against SARS-CoV-2 can enter cells readily and inhibit viral replication in a sequence-specific, dose-responsive and non-toxic manner. Considering the considerable *in vivo* antiviral efficacy demonstrated by PPMO against several respiratory viruses in previous studies [5–8], further development of the 5’END- and TRS-PPMO appears warranted.

## Transparency declaration

The authors have no conflicts of interest.

## References

1. Moulton, H.M. and J.D. Moulton, Morpholinos and their peptide conjugates: therapeutic promise and challenge for Duchenne muscular dystrophy. Biochim Biophys Acta, 2010. 1798(12): p. 2296–303.

2. Moulton, H.M., and J.D. Moulton, Antisense Morpholino Oligomers and Their Peptide Conjugates, in Therapeutic Oligonucleotides, J. Kurreck, Editor. 2008, Royal Society of Chemistry: London, England. p. 43–79.

3. Abes, S., et al., Vectorization of morpholino oligomers by the (R-Ahx-R)(4)peptide allows efficient splicing correction in the absence of endosomolytic agents. J Control Release, 2006. 116(3): p. 304–13.

4. Stein, D.A., Inhibition of RNA virus infections with peptide-conjugated morpholino oligomers. Curr Pharm Des, 2008. 14(25): p. 2619–34.

5. Gabriel, G., et al., Morpholino oligomers targeting the PB1 and NP genes enhance the survival of mice infected with highly pathogenic influenza A H7N7 virus. J Gen Virol, 2008. 89(Pt 4): p. 939–48.

6. Lupfer, C., et al., Inhibition of influenza A H3N8 virus infections in mice by morpholino oligomers. Archives of virology, 2008. 153(5): p. 929–37.

7. Lai, S.H., et al., Inhibition of respiratory syncytial virus infections with morpholino oligomers in cell cultures and in mice. Mol Ther, 2008. 16(6): p. 1120–8.

8. Opriessnig, T., et al., Inhibition of porcine reproductive and respiratory syndrome virus infection in piglets by a peptide-conjugated morpholino oligomer. Antiviral Res, 2011. 91(1): p. 36–42.

9. Masters, P.S., and S. Perlman, Coronaviridae, in Fields virology, D.M. Knipe, and P.M. Howley, Editor. 2013, Lippincott Williams & Wilkins: Philadelphia, PA. p. 825–858.

10. Burrer, R., et al., Antiviral effects of antisense morpholino oligomers in murine coronavirus infection models. J Virol, 2007. 81(11): p. 5637–48.

11. Neuman, B.W., et al., Antisense morpholino-oligomers directed against the 5’ end of the genome inhibit coronavirus proliferation and growth. J Virol, 2004. 78(11): p. 5891–9.

12. Neuman, B.W., et al., Inhibition, escape, and attenuated growth of severe acute respiratory syndrome coronavirus treated with antisense morpholino oligomers. J Virol, 2005. 79(15): p. 9665–76.

13. Munster, V.J., et al., Respiratory disease in rhesus macaques inoculated with SARS-CoV- 2. Nature, 2020.

14. van den Born, E., et al., Antiviral activity of morpholino oligomers designed to block various aspects of Equine arteritis virus amplification in cell culture. J Gen Virol, 2005. 86(Pt 11): p. 3081–90.

15. Zhang, Y.J., et al., Suppression of porcine reproductive and respiratory syndrome virus replication by morpholino antisense oligomers. Vet Microbiol, 2006. 117(2-4): p. 117–29.

16. Sawicki, S.G., D.L. Sawicki, and S.G. Siddell, A contemporary view of coronavirus transcription. J Virol, 2007. 81(1): p. 20–9.

17. Pasternak, A.O., W.J. Spaan, and E.J. Snijder, Nidovirus transcription: how to make sense…? J Gen Virol, 2006. 87(Pt 6): p. 1403–21.

18. Yang, D. and J.L. Leibowitz, The structure and functions of coronavirus genomic 3’ and 5’ ends. Virus Res, 2015. 206: p. 120–33.

19. Khailany, R.A., M. Safdar, and M. Ozaslan, Genomic characterization of a novel SARS-CoV-2. Gene Rep, 2020: p. 100682.

20. Ge, Q., et al., Inhibition of multiple subtypes of influenza a virus in cell cultures with morpholino oligomers. Antimicrob Agents Chemother, 2006. 50(11): p. 3724–33.

